# Comprehensive in-vivo secondary structure of the SARS-CoV-2 genome reveals novel regulatory motifs and mechanisms

**DOI:** 10.1101/2020.07.10.197079

**Authors:** Nicholas C. Huston, Han Wan, Rafael de Cesaris Araujo Tavares, Craig Wilen, Anna Marie Pyle

## Abstract

SARS-CoV-2 is the positive-sense RNA virus that causes COVID-19, a disease that has triggered a major human health and economic crisis. The genome of SARS-CoV-2 is unique among viral RNAs in its vast potential to form stable RNA structures and yet, as much as 97% of its 30 kilobases have not been structurally explored in the context of a viral infection. Our limited knowledge of SARS-CoV-2 genomic architecture is a fundamental limitation to both our mechanistic understanding of coronavirus life cycle and the development of COVID-19 RNA-based therapeutics. Here, we apply a novel long amplicon strategy to determine for the first time the secondary structure of the SARS-CoV-2 RNA genome probed in infected cells. In addition to the conserved structural motifs at the viral termini, we report new structural features like a conformationally flexible programmed ribosomal frameshifting pseudoknot, and a host of novel RNA structures, each of which highlights the importance of studying viral structures in their native genomic context. Our in-depth structural analysis reveals extensive networks of well-folded RNA structures throughout Orf1ab and reveals new aspects of SARS-CoV-2 genome architecture that distinguish it from other single-stranded, positive-sense RNA viruses. Evolutionary analysis of RNA structures in SARS-CoV-2 shows that several features of its genomic structure are conserved across beta coronaviruses and we pinpoint individual regions of well-folded RNA structure that merit downstream functional analysis. The native, complete secondary structure of SAR-CoV-2 presented here is a roadmap that will facilitate focused studies on mechanisms of replication, translation and packaging, and guide the identification of new RNA drug targets against COVID-19.

## INTRODUCTION

Severe acute respiratory syndrome related coronavirus 2 (SARS-CoV2), which is responsible for the current global pandemic(Zhu et al., 2020), is a positive strand RNA virus in the genus *β-coronavirus*. To date, the outbreak of SARS-CoV2 has infected at least 12 million people globally, causing great economic loss and posing an ongoing public health threat(Dong et al., 2020). Included in the β-coronavirus genus are two related viruses, SARS-CoV and Middle East respiratory syndrome coronavirus (MERS-CoV), that caused global outbreaks in 2003 and 2012, respectively(de Wit et al., 2016). Despite the continued risk posed by β-coronaviruses, mechanistic studies of the family are limited and to date no effective antivirals or vaccines exist, highlighting the need for research that facilitates the development of therapeutics. With most research efforts focusing on viral proteins (Lan et al., 2020a, Yin et al., 2020, Wan et al., 2020), little is known about the viral RNA genome, especially its structural content. This is a major gap in our understanding because RNA structural elements in positive strand viruses play central roles in regulating all aspects of replication, translation, packaging and host defense (McMullan et al., 2007, Fricke et al., 2015, Pirakitikulr et al., 2016, Clyde and Harris, 2006, MacFadden et al., 2018), so an understanding of their location and function is critical for mechanistic understanding and strategies for viral control(Barrows et al., 2018, Adams et al., 2017). Given the success of antimicrobials targeted against conserved RNA structural elements in other pathogen genomes(Warner et al., 2018, Fedorova et al., 2018), there is an urgent, unmet need to elucidate the genome architecture of SARS-CoV-2.

Like other coronaviruses, the genome of SARS-CoV-2 is incredibly large (Maier et al., 2015, Zhu et al., 2020). Two open reading frames (ORF) for viral nonstructural proteins (Nsp) and 9 small ORFs that encode the structural proteins and a number of accessory genes comprise a ∼30kb genome(Kim et al., 2020). These ORFs are flanked on either side by a 5’ and 3’UTR that have been shown in other coronaviruses to possess conserved RNA structures with important functional roles in the viral life cycle(Yang and Leibowitz, 2015). Studies in Murine Hepatitis Virus (MHV) and Bovine Coronavirus (BCoV) suggest that the 5’ viral termini folds in to 6 stems (SL1-SL6) that play roles in sgRNA synthesis and viral replication(Madhugiri et al., 2018, Chen and Olsthoorn, 2010). In the 3’UTR, a pseudoknot and a bulged stem loop (BSL) are essential for sgRNA synthesis in MHV(Zust et al., 2008).

One of the best-studied functional RNA elements in β-coronavirus genomes is the programmed ribosomal frameshifting pseudoknot (PRF) that sits at the boundary between Orf1a and Orf1ab(Plant and Dinman, 2008). The PRF, found in all coronaviruses, induces a −1 ribosomal frameshift that allows for bypassing of the Orf1a stop codon and production of the orf1ab polyprotein, which includes the viral replicase(Plant et al., 2005). Reporter assays using a truncated PRF construct showed that programmed frameshifting occurs ∼25% of the time in SARS-CoV(Kelly et al., 2020), and that it is crucial for sgRNA synthesis(Plant et al., 2013). Extensive mutational analysis has revealed a three-stemmed pseudoknot structure for the SARS-CoV PRF(Plant et al., 2005). However, neither the mechanism of frameshifting regulation nor the three-stem pseudoknot PRF conformation has been validated in cells.

While recent computational studies suggest the 5’UTR, 3’UTR, and PRF functional elements are conserved in the SARS-CoV-2 genome(Rangan et al., 2020, Andrews et al., 2020), these regions account for a vanishingly small fraction of the total nucleotide content. Studies of other positive-sense viral RNA genomes such as Hepatitis C virus (HCV) and Human Immunodeficiency Virus (HIV) have revealed extensive networks of regulatory RNA structures contained within viral ORFs(Siegfried et al., 2014, Pirakitikulr et al., 2016, Friebe and Bartenschlager, 2009, Li et al., 2018, You et al., 2004) which direct critical aspects of viral function. It is therefore of crucial importance to assess and characterize the structural features of the SARS-CoV-2 ORF, as elucidation of structural motifs will improve our understanding of all coronaviruses and facilitate development of antiviral therapies for the entire family.

Recent advances in high-throughput structure probing methods (SHAPE-MaP, DMS-MaP) have greatly facilitated the structural studies of long viral RNAs (Siegfried et al., 2014, Zubradt et al., 2017). Recently, Manfredonia et al. performed full-length SHAPE-MaP analysis on *ex vivo* extracted and refolded SARS-CoV-2 RNA(Manfredonia et al., 2020). However, structural studies on both viral and messenger RNA have highlighted the importance of probing RNAs in their natural cellular context(Simon et al., 2019, Rouskin et al., 2014). Lan et al performed full-length *in-vivo* DMS-MaPseq on SARS-CoV2 infected cells(Lan et al., 2020b), but as DMS only reports on A and C nucleotides, the data coverage is necessarily sparse. While both studies reveal important features of the structural content in the SARS-CoV-2 genome and its evolutionary conservation, to date no work has been published that captures information for every single nucleotide in an *in-vivo* context.

Here, we report for the first time the complete secondary structure of SARS-CoV-2 RNA genome using in SHAPE-MaP data obtained in living cells. We deploy a novel long amplicon method readily adapted to other long viral RNAs made possible by the highly processive reverse transcriptase MarathonRT (Guo et al., 2020). The resulting genomic secondary structure map reveals functional motifs at the viral termini that are structurally homologous to other coronaviruses, thereby fast-tracking our understanding of the SARS-CoV-2 life cycle. We reveal conformational variability in the PRF, highlighting the importance of studying viral structures in their native genomic context and underscoring their dynamic nature. We also uncover elaborate networks of well-folded RNA structures dispersed across Orf1ab, and we reveal features of the SARS-CoV-2 genome architecture that distinguish it from other single-stranded, positive-sense RNA viruses. The analysis reveals large RNA structures within the ORF that may ultimately prove to be as important for viral function as the PRF. Evolutionary analysis of the full-length SARS-CoV-2 structure suggests that, not only do its architectural features appear to be conserved across the β-coronavirus family, but individual regions of well-folded RNA may be as well. Our work reveals the unique genomic architecture of SARS-CoV-2 in infected cells, points to important viral strategies for infection and persistence, and identifies potential drug targets. The full-length structure model we present here thus serves as an invaluable roadmap for future studies on SARS-CoV-2 and other coronaviruses that emerge in the future.

## MATERIALS & METHODS

### Cell Culture and SARS-CoV-2 Infection

VeroE6 cells were cultured in Dulbecco’s Modified Eagle Medium (DMEM) with 10% heat-inactivated fetal bovine serum (FBS). Approximately 5×10^6^ cells were plated in each of four T150 tissue culture treated flasks. The following day media was removed and 10^5^ PFU in 4mL of media of SARS-CoV-2 isolate USA-WA1/2020 (BEI Resources #NR-52281) was added to each flask. Virus was adsorbed for 1 hour at 37°C and then 16mL of fresh media was added to each flask.

### RNA Probing and Purification

Four days post-infection (dpi), the supernatant was aspirated from each flask, cells were washed with 10mL of cold PBS-/- and then dislodged in 10ml PBS-/- with a cell scraper. The contents were collected and centrifuged at 450g x 5 min at 4°C. The supernatant was removed and the cell pellet was resuspended into 2ml of PBS-/- with 200μl DMSO or 2ml PBS with 200μl of 2M NAI (final concentration = 200mM). Cells were incubated for 10 minutes at room temperature followed by addition of 6mL of Trizol. RNA was extracted with the addition of 1.2mL of chloroform. The aqueous phase was transferred to a new tube, followed by the addition of 12mL of 100% EtOH (70% final) and precipitated overnight at −20°C. RNA was resuspended in 1xME buffer and purified using the Qiagen RNeasy kit according to the manufacturer’s protocol. RNA was eluted in 1xME buffer.

### Tiled-Amplicon Design

Leveraging the extreme processivity of MarathonRT, a highly processive group II intron-encoded RT(2), we designed fifteen 2000nt amplicon and a single 1300nt amplicons tiled across the SARS-CoV-2 genome for full sequencing coverage. Adjacent amplicons were designed with a 100nt overlap to ensure data is collected for regions otherwise masked by primer binding. Primers for reverse transcription (RT) were designed using the OligoWalk tool(Lu and Mathews, 2008) to avoid highly-structured primers and highly-structured regions of the SARS-CoV-2 genome. Forward and reverse primer sets were designed for an optimal T_m_ of 58°C. Reverse primers were inset 3nt from the 5’end of the RT primer to enhance specificity of the PCR reaction.

### Reverse Transcription with MarathonRT

MarathonRT purification was performed as described in (Guo et al., 2020). For each amplicon, 500ng of total cellular RNA was mixed with 1μL of the corresponding 1μM RT primer. Gene-specific primers used for RT are listed in **Table S2**. Primers were annealed at 65°C for 5min then cooled to room temperature, followed by addition of 8μL of 2.5x MarathonRT SHAPE-Map Buffer (125mM 1M Tris-HCl pH 7.5, 500mM KCl, 12.5mM DTT, 1.25mM dNTPs, 2.5mM Mn^2+^), 4μL of 100% glycerol, and 0.5μL of MarathonRT. RT reactions were incubated at 42°C for 3 hours. 1μL 3M NaOH was added to each reaction and incubated at 95°C for 5min to degrade the RNA, followed by the addition of 1μL 3M HCl to neutralize the reaction. cDNA was purified using AmpureXP beads (Cat. No. A63880) according to manufacturer’s protocol and a 1.8x bead-to-sample ratio. Purified cDNA was eluted in 10μL nuclease-free water.

### SHAPE-MaP Library Construction

Amplicons tiling the SARS-CoV-2 genome were generated using NEBNext UltraII Q5 MasterMix (Cat. No. M0544L), gene-specific forward and reverse PCR primers, and 5μL of purified cDNA. Gene-specific primers used for PCR are listed in **Table S3**. Touchdown cycling PCR conditions were used to enhance PCR specificity (68-58°C annealing temperature gradient). PCR reaction products were purified with Monarch DNA Clean-up Kits (NEB) with a binding buffer:sample ratio of 2:1 to remove products smaller than 2kb. PCR products were visualized on 0.8% agarose gels to confirm production of correctly sized amplicons. Amplicons were diluted to 0.2ng/uL and then pooled into two odd and two even amplicon pools for downstream library preparation. Sequencing libraries were generated using a NexteraXT DNA Library Preparation Kit (Illumina) according to manufacturer’s protocol, but with 1/5^th^ the recommended volume. Libraries were quantified using a Qubit (Life Technologies) and a BioAnalyzer (Agilent). Amplicon pools were recombined and sequenced on a NextSeq 500/550 platform using a 150 cycle mid-output kit.

### Structure Prediction

All libraries were analyzed using ShapeMapper 2(Busan and Weeks, 2018), aligning reads to SARS-CoV-2 genome (accession number: MN908947). Mutation rates between NAI-modified and unmodified samples were tested for significance using the equal variance t-test. Using reactivities output from ShapeMapper, ShapeKnots(Hajdin et al., 2013) was used to determine whether two previously reported pseudoknots contained in the SARS-CoV-2 genome were predicted with experimental SHAPE constraints. The two pseudoknots tested were the programmed ribosomal frameshifting element that exists at the Orf1a/b boundary, and a pseudoknot in the 3’UTR that was identified in the MHV and B-CoV genomes (references). We analyzed all 500nt windows separated by a 100nt slide that contained each of the putative pseudoknots to determine if the pseudoknot was successfully predicted.

SuperFold (Smola et al., 2015b) was used to generate a consensus structure prediction for the entire SARS-CoV-2 genome using SHAPE reactivities obtained from biological replicate 1 as constraints. We imposed a maximum pairing distance of 500nt. As our data only supported formation of the pseudoknot contained in the programmed ribosomal frameshifting element, only this pseudoknot was forced in this prediction. All structures output from the SuperFold prediction were visualized and drawn using StructureEditor, a tool in the RNAStructure software suite(Reuter and Mathews, 2010).

Base-pairing distances were calculated from .ct structure files output from SuperFold full-length SARS-CoV-2 consensus predictions, and compared to previously published, publically available full-length genome structures for Dengue and Hepatitis C Virus generated with SHAPE constraints, a max-pairing distance of 500nt, and the SuperFold pipeline (Mauger et al., 2015, Dethoff et al., 2018).

### Identification of Well-Folded Regions

Two data signatures were used to identify well-folded regions: The first is the SHAPE reactivity data generated with the SHAPE-MaP workflow and the ShapeMapper analysis tool(Busan and Weeks, 2018). The second is the Shannon entropy calculated from base-pairing probabilities determined during the SuperFold partition function calculation(Smola et al., 2015b). Two replicate data sets were used, including separate SuperFold predictions.

Local median SHAPE reactivity and Shannon Entropy were calculated in 55nt sliding windows. The global median SHAPE reactivity or Shannon Entropy were subtracted from calculated values to aid in data visualization. Regions with local SHAPE and Shannon Entropy signals 1) below the global median 2) for stretches longer than 40 nucleotides 3) that appear in both replicate data sets were considered well-folded. Disruptions, or regions where local SHAPE or Shannon Entropy rose above the global median, are not considered to disqualify well-folded regions if they extended for less than 40 nucleotides. Arc plots generated from each replicate consensus structure predication were compared for regions that meet sorting criteria described above in order to ensure agreement between secondary structure models generated from each replicate SHAPE-MaP dataset.

Base-pairing distances of well-folded regions were calculated from .ct structure files output from SuperFold consensus predictions, and compared to previously published, publicly available structures for well-folded regions of the HIV genome generated with SHAPE constraints, a max-pairing distance of 500nt, and the SuperFold pipeline (Siegfried et al., 2014).

### Multiple sequence alignment

To analyze evolutionary support for our consensus secondary structure prediction of the SARS-CoV-2 genome, we generated two codon-based multiple sequence alignments (MSA) for Orf1a and Orf1b constructed from genomes of closely related viral species (Douzery EJP,2018). All sequences were chosen based on a phylogenetic study of SARS-CoV-2 (Ceraolo and Giorgi, 2020). All sequences referenced below were downloaded from the NCBI Taxonomy browser(Benson et al., 2018).

A sarbecovirus MSA was generated using SARS-CoV-2 isolate Wuhan-Hu-1 (MN908947.3), four bat coronaviruses (MG772934.1, JX993987.1, DQ022305.2, DQ648857.1), and five human SARS coronaviruses (AY515512.1, AY274119.3, NC_004718.3, GU553363.1, DQ182595.1).

We also generated an “All β-coronavirus Alignment” using the sarbecovirus sequences described above in addition to four MERS-CoV sequences (MK129253, KP209307, MF598594, MG987420), one HKU-4 sequence (MH002337), three HKU-5 sequence (MH002342, NC009020, MH002341), four HKU1 sequences (KY674942, KF686343, AY597011, DQ415903), three murine hepatitis virus sequences (AY700211, AF208067, AB551247), three human coronavirus OC43 sequences (AY585229, NC006213, MN026164), two bovine coronavirus sequences (KU558922, KU558923), and one camel coronavirus sequence (MN514966).

The orf1a and orf1b region were extracted from the full-length sequences based on the GenBank annotation. Separate codon alignments for both Orf1a and orf1b were generated using MACSE v2.0.3(Ranwez et al., 2018) and default parameters (*-prog alignSequences*).

### Synonymous mutation rate analysis

All codon alignments were visualized and edited using Jalview v 2.11.0(Waterhouse et al., 2009). Synonymous mutation rates for each codon were estimated using the phylogenetic-based parametric maximum likelihood (FUBAR) method(Murrell et al., 2013). Each codon was categorized as base-paired or unpaired depending on strandedness of the nucleotide at the third position of each codon in our SARS-CoV-2 consensus structure model(Dethoff et al., 2018). The significance of synonymous mutation rates between single- and double-stranded regions was determined using two-tailed, equal variance *t*-test.

### Covariation analysis

Covariation calculation and visualization was performed using R-chie(Lai et al., 2012). The Sarbecovirus codon alignment described above was used for covariation analysis. Identification of base-pairs with statistically significant evidence of covariation was performed on individual structures using R-Scape (version 0.2.1)(Rivas et al., 2017) with the RAFSp statistics by using the “--RAFSp” flag(default E-value:0.05)(Tavares et al., 2019).

### Data Availability

All ShapeMapper outputs, secondary structure files, and multiple sequence alignments use in this work are available at the GitHub repository: https://github.com/pylelab/SARS-CoV-2_SHAPE_MaP_structure

## RESULTS

### *In-vivo* SHAPE-MaP workflow yields high quality data suitable for structure prediction

To study the SARS-CoV-2 structure in the context of infected cells, the SARS-CoV-2 isolate USA-WA1/2020, isolated from an oropharyngeal swab from a patient who had returned to the United States from China and developed clinical disease, was used to infect VeroE6 cells (BEI Resources #NR-52281). Infection was allowed to proceed for four days, at which point cells were collected and treated with either NAI or DMSO. RNA was then extracted and purified. To generate sequencing libraries, 2000 nucleotide (nt) overlapping amplicons were tiled across the entire SARS-CoV-2 genome (**Fig. 1A**). Importantly, this approach is made possible by the utilization of the ultra-high processive reverse transcriptase MarathonRT, which encodes NAI adducts as cDNA mutations during long-amplicon SHAPE-MaP library construction(Guo et al., 2020). Specifically, gene-specific primers were used to prime reverse transcription in the presence of manganese, followed by amplicon PCR with gene-specific primers and cycling conditions designed to enhance specificity(Korbie and Mattick, 2008). Gel electrophoresis confirmed successful amplification of all 16 amplicons (data not shown). Sequencing of SHAPE-MaP libraries was performed using the Illumina NextSeq 500/550 platform.

**Figure 1.**
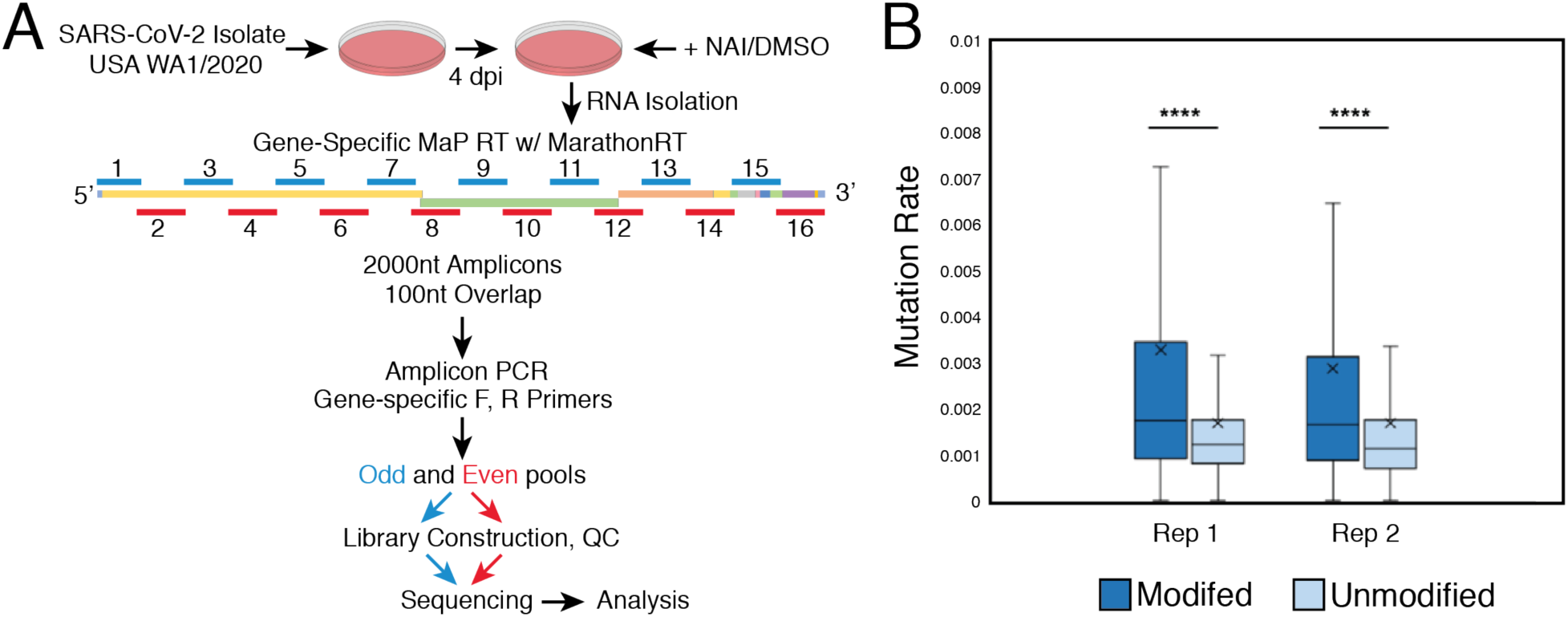
Tiled-amplicon *in-vivo* SHAPE-MaP workflow yields high quality data for SARS-CoV-2 structure prediction. **A)** Workflow of *in-vivo* SHAPE-MaP probing of full-length SARS-CoV-2 genomic RNA. **B)** Mutation rates for two biological replicates confirm genomic RNA was successfully modified with NAI electrophile. The boxes represent the interquartile range (IQR) of each data-set, with the median value indicated by a line, average value indicated by a “x”. Tukey-style whiskers extend 1.5 x IQR beyond each box. Values outside this range are not shown. ****p<0.0001 by equal variance unpaired student t test.

After generating two independent biological replicates, the resulting sequencing data were analyzed using the ShapeMapper pipeline(Smola et al., 2015b). Comprehensive datasets were obtained, with median effective read depth > 70,000x and effective reactivity data for 99.7% (29813/29903) of nucleotides in the SARS-CoV-2 genome in both replicate experiments. To check the SHAPE-MaP data quality, we analyzed the relative mutation rates of NAI-treated and DMSO-treated RNA samples, revealing a significant elevation of mutation rates for NAI-treated samples (**Fig. 1B**, p-value < 0.0001). This confirms that the full-length SARS-CoV-2 RNA was successfully modified *in-vivo* and that these modifications were encoded as cDNA mutations.

To understand the relative SHAPE reactivity agreement within local regions of the genome, we calculated Pearson correlation coefficients between two biological replicates. The Pearson’s correlation across the entire span of Orf1ab is 0.62, consistent with those previously reported for reactivities calculated from *in-vivo* modified RNAs of this size(Smola et al., 2016). Across the sub genomic RNA ORFs, the Pearson ‘s correlation is poor. We believe this reflects the fact that Amplicons 13, 14, 15, and 16 will amplify both full-length *and* sub-genomic RNAs, and the difference in context will result in different secondary structures(Tavares et al., 2020). For this reason, despite the fact all data have been obtained globally, subsequent discrete structural analysis will focus on shared features of the viral termini and the Orf1ab region.

### *De novo* structure prediction on full-length SARS-CoV-2 RNA identifies conserved functional elements at the 5’ and 3’ genomic termini

We performed secondary structure prediction with the SuperFold pipeline(Smola et al., 2015b), using the *in-vivo* SHAPE reactivities to generate an experimentally constrained consensus secondary structure prediction for the entire SARS-CoV-2 genome. As an extensive body of research has elucidated structured RNA elements at the 5’ and 3’ viral termini as well as the Orf1ab boundary, with conserved functions across β-coronaviruses, we first examined these regions from our consensus prediction to determine whether they were stably folded and well-determined in the SARS-CoV-2 genome.

The 5’ genomic terminus includes seven regions that have been identified and studied in other coronaviruses (Reviewed in (Yang and Leibowitz, 2015)). While sequence conservation suggested that these elements might be conserved in SARS-CoV-2, our consensus structure prediction shows this to be the case, and we derived a specific experimentally-determined structure for this section of the genome. The in-vivo SHAPE reactivity data correspond well with the resulting structural model (**Fig 2A, inset**) and the low overall Shannon entropy values in this region (determined from base pair probability calculation during the SuperFold prediction pipeline(Smola et al., 2015b)) support a well-determined structure for the 5’ genomic terminus (median_Nuc(1-400)_ = 2.7×10^−5^; global median = 0.022).

**Figure 2.**
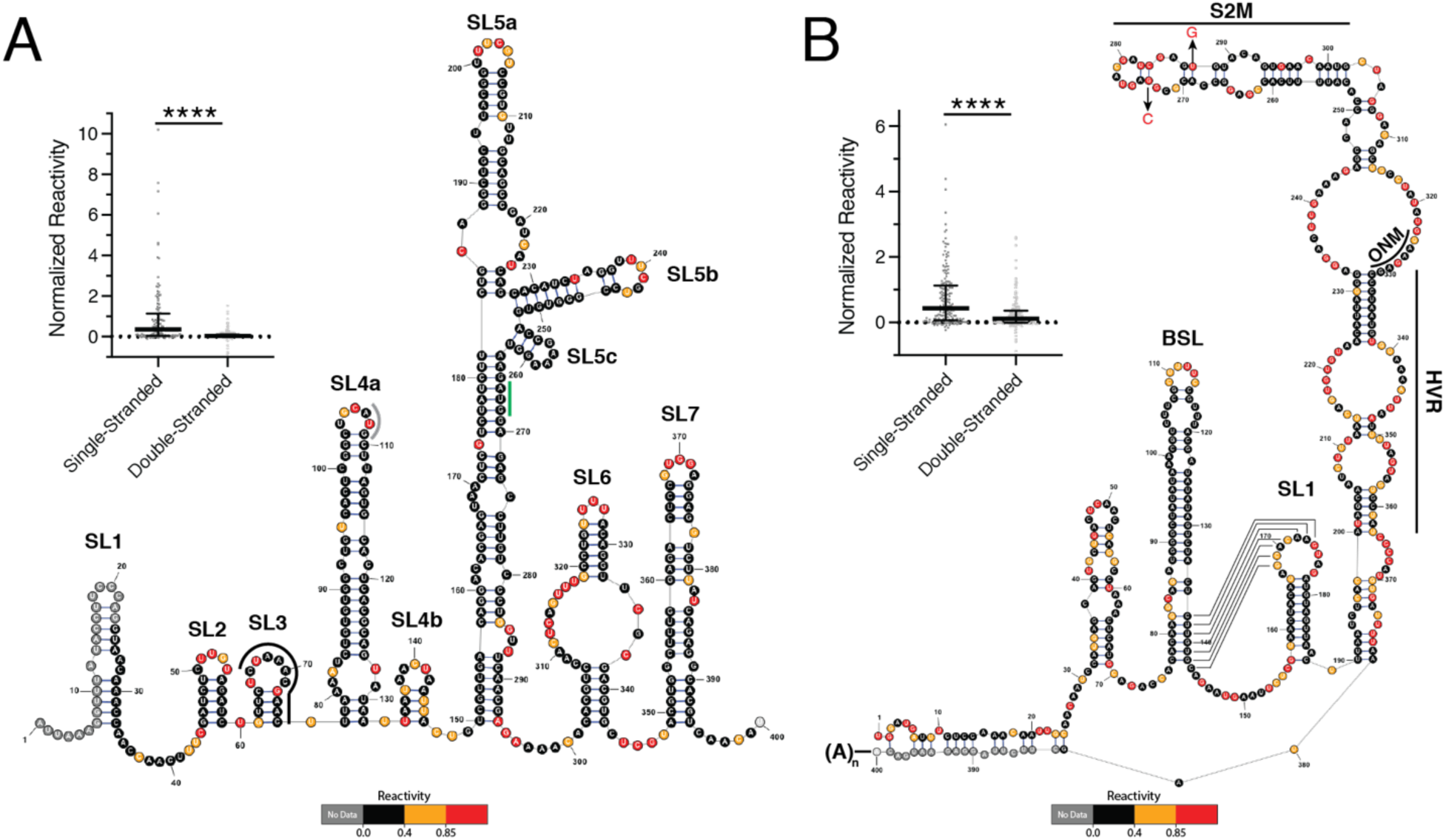
*De novo* full-length structure prediction of SARS-CoV-2 genomic RNA identifies conserved functional elements at the 5’ and 3’ viral termini. **A)** Consensus structure prediction for the 5’ terminus of SARS-CoV-2, colored by SHAPE Reactivity. Functional domains are labeled, including TRS sequence, start codon of uORF, and start codon of Orf1a (indicated by black, grey, and green lines, respectively). Inset – mapping of SHAPE reactivity data to single- and double-stranded regions, data are plotted with a line indicating the median, and whiskers indicating the standard deviation **B)** Structure prediction for the 3’ terminus of SARS-CoV-2, colored by SHAPE reactivity. Functional domains are labeled. The putative pseudoknot is indicated by solid black lines. Locations of the octanucleotide motif (ONM), hypervariable region (HVR) and S2M are indicated by black lines. Inset – mapping of SHAPE reactivity to single- and double-stranded regions. Data are plotted with a line indicating the median, and whiskers indicating the standard deviation ****p<0.0001 by equal variance unpaired student t test.

Individual features that typify coronavirus structures are evident in the secondary structure of the SARS-CoV-2 5’-UTR with good SHAPE reactivity agreement (**Fig 2A**, inset). For example, we observe a bipartite domain architecture for SL1, which was previously reported to play a role in coronavirus replication(Li et al., 2008) (**Fig 2A**, labeled SL1). Similarity between SL1 structures reported for other coronaviruses and the experimentally-determined structure reported here for SARS-CoV-2 suggests that SL1 plays a similar role in SARS-CoV-2 life cycle.

Structural studies of SARS-CoV SL2 have shown that the SL2 pentaloop is stacked atop a 5-bp stem. In addition, the pentaloop adopts a canonical CUYG fold in which the uracil is flipped out, resulting in an architecture that is important for sgRNA synthesis(Lee et al., 2011). Our experimentally-determined structure of SL2 from SARS-CoV-2 shows that it adopts exactly the same RNA fold (**Fig 2A**, labeled SL2), again suggesting that it plays the same functional role in the SARS-CoV-2 life cycle.

The transcription regulatory sequence (TRS) is a conserved feature of β-coronaviruses and it is required for sgRNA production(Yang and Leibowitz, 2015). The SARS-CoV leader TRS is predicted *in silico* to be in stem loop 3 (SL3), with nucleotides exposed in its loop and base-paired in the stem(Chen and Olsthoorn, 2010). The primary sequence of the SARS-CoV leader TRS is absolutely conserved between SARS-CoV and SARS-CoV-2(Chen and Olsthoorn, 2010) (5’-ACGAAC-3’). Importantly, our consensus prediction shows the SARS-CoV-2 leader TRS is also found in SL3, with a similar structural organization as reported for other viruses (**Fig 2A**, labeled SL3, TRS indicated with solid black line).

The SL4 of SARS-CoV-2 adopts a bipartite domain structure (**Fig 2A**, labeled SL4a, SL4b) similar to that reported for MHV(Kang et al., 2006, Yang et al., 2015). Importantly, the AUG of the predicted upstream ORF(uORF), which is phylogenetically conserved among β-coronaviruses(Raman et al., 2003), is found in the top-most stem loop of SL4a, meaning it would be accessible for recognition by a scanning ribosome (**Fig 2A**, uORF start codon indicated with solid grey line).

As predicted across coronaviruses(Chen and Olsthoorn, 2010), the trifurcated stem at the top of SL5 is observed in the experimentally-determined structure of SARS-CoV-2 (**Fig 2A**, labeled SL5A-C). This includes UUCGU pentaloop motifs in SL5A and SL5B, and a GNRA tetraloop in SLC. Previous reports suggest this may represent a packaging signal for GroupIIB CoVs(Chen and Olsthoorn, 2010).

SL6 and SL7 are predicted in the SARS-CoV-2 structure, and the in-vivo SHAPE data agree strongly support the existence of these stems (**Fig 2A**, labeled SL6 and SL7). However, functional evidence for SL6 and SL7 is lacking in the literature for any coronavirus.

The 3’ genomic terminus includes three well-studied stems, including the bulged-stem loop (BSL), Stem Loop 1 (SLI), and a long-bulge stem that includes the hypervariable-region (HVR), the S2M domain, the octanucleotide motif (ONM) subdomains, and a pseudoknot (Reviewed in (Yang and Leibowitz, 2015)). The consensus structure recapitulates the secondary structure of all the three stems with good SHAPE reactivity agreement (**Fig 2B, inset**) and overall low Shannon entropy (median_Nuc(29,472-29,870)_ = 0.016; global median = 0.022). While the BSL is well determined in our structure, the low reactivity for bulged nucleotides suggests the possibility of protein binding-partners (**Fig 2B**, labeled BSL).

A pseudoknot structure is proposed to exist between the base of the BSL stem loop and the loop of SL1 in coronaviruses(Yang and Leibowitz, 2015). While pseudoknot formation is mutually exclusive with the base of the BSL, studies in MHV have suggested that both structures contribute to viral replication and the mutually exclusive structures are thought to function as a molecular switch in different steps of RNA synthesis(Goebel et al., 2004). However, our *in-vivo* determined secondary structure is inconsistent with formation of the pseudoknot (**Fig 2B**, putative base-pairing interactions indicated by black lines). The low SHAPE reactivities for the nucleotides at the base of the BSL support formation of the extended BSL stem, while high-reactivities of the nucleotides in the loop of SLI indicate that it is highly accessible. Using the SHAPEKnots program for robust predication of pseudoknots (implemented in RNA structure v5.8(Hajdin et al., 2013)), we found that a pseudoknot is never predicted in three 500nt windows that cover the pseudoknotted region. Taken together, our data strongly support the extended BSL conformation, indicating it is probably the dominant conformation *in-vivo*.

The third stem in the 3’ UTR includes three sub-domains. The HVR, so-named because it is poorly conserved across group II coronaviruses(Goebel et al., 2007), is predicted to be mostly single-stranded in our secondary structure, and the high reactivities across the span of this region lends strong experimental support for an unstructured region (**Fig. 2B**, region indicated with solid black line and labeled HVR). The fact that this region is highly unstructured may also explain why it has been experimentally demonstrated to tolerate numerous deletions, rearrangements, and point mutations in MHV(Goebel et al., 2007).

The S2M region is contained within the apical part of the third stem. We observe that the first three helices of S2M from SARS-CoV-2 exactly match the crystal structure determined for S2M from SARS-CoV (Robertson et al., 2005). However, our in-vivo structure deviates significantly at the top of the stem, with bases that are highly reactive (**Fig. 2B**, region indicated with solid black line and labeled S2M). It is possible that the SARS-CoV-2 S2M folds into a unique S2M conformation despite differing by only a two bases (**Fig. 2B**, base-changes indicated by arrows; SARS-CoV base identity shown in red). Indeed, as both single-nucleotide changes are transversions, any base-pairing interaction involving these nucleotides in the SARS-CoV S2M structure could not be maintained in SARS-CoV-2. Alternatively, this site could interact with factors *in-vivo* that are not captured in the crystallographic study.

The ONM is predicted at the central bulge between the S2M region and the HVR region. The sequence is absolutely conserved across β-coronavirus(Goebel et al., 2007), but no functional significance has yet been shown. In our consensus structure, it is single-stranded (**Fig. 2B**, ONM indicated with solid black line and labeled ONM).

Finally, we predict a completely different structure for the downstream terminal stem in the viral 3’UTR region (adjacent to the poly-A tail) than previously reported for other coronaviruses(Zust et al., 2008). However, our structure prediction in this region is not highly accurate because of proximity of the primer binding site. That said, the putative stem is predicted to have high Shannon entropy (median_Nuc(29472-29495,29853-29870)_ = 0.2154; global median = 0.022), suggesting that it is not a well-ordered structure in the cellular environment.

### Structure prediction of the programmed ribosomal frame-shifting element reveals conformational flexibility

One of the most well-studied RNA structures in the coronavirus coding region is the programmed frame-shifting pseudoknot (PRF). It is located between orf1a and orf1b and plays an important role in inducing a −1 frameshift in a translating ribosome, resulting in the synthesis of the polyprotein ab, which includes the SARS-CoV-2 replicase (Plant and Dinman, 2008).

The PRF element previously characterized in SARS-CoV is proposed to contain three parts: an attenuator stem loop, a conserved heptanucleotide “slippery” sequence, and a H-type pseudoknot (Plant and Dinman, 2008). We performed SHAPEKnots(Hajdin et al., 2013) over four 500nt windows that cover the pseudoknotted region in the SARS-CoV-2 genome to check if the PRF pseudoknot can be discerned from our vivo SHAPE data. We found that the pseudoknot is successfully predicted in 3 out of 4 windows generated by ShapeKnots. Moreover, the nucleotides predicted to be involved in the pseudoknotted helix have low SHAPE-reactivity (**Fig. 3A**, pseudoknot base-pairs indicated with red lines). Our *in-vivo* SHAPE data therefore strongly support the formation of the pseudoknotted helix, and the frame-shifting pseudoknot was thereafter included as a hard constraint during secondary structure prediction.

**Figure 3.**
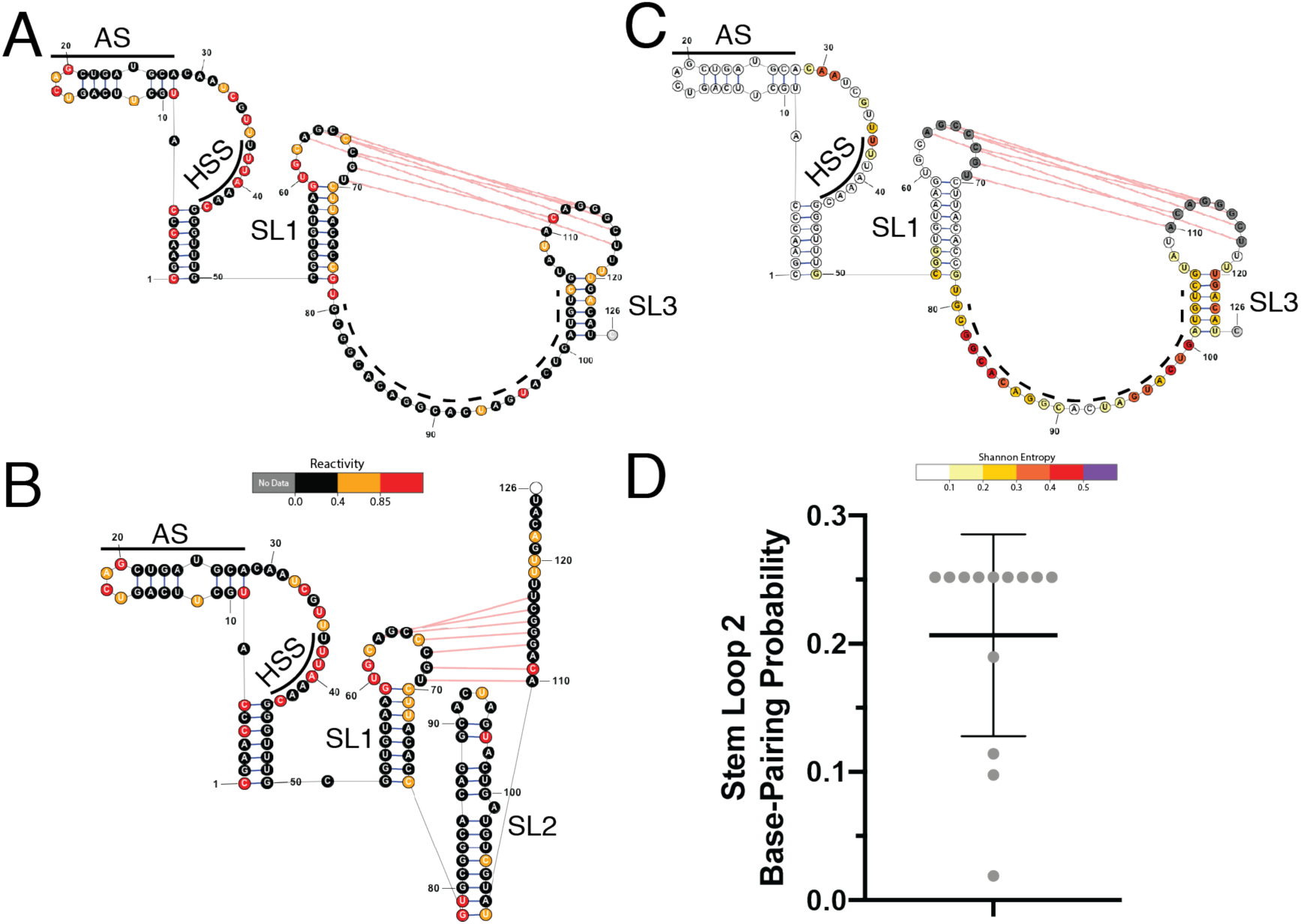
Structure prediction of the programmed ribosomal frame-shifting (PRF) element suggests conformational variability of Stem Loop 2. **A)** Dominant PRF structural architecture, predicted by SuperFold, colored by relative SHAPE Reactivity from this study. AS = Attenuator Stem; HSS = Heptanucleotide Slippery Sequence; SL1 = Stem Loop 1; dotted line indicates region reported to form stem loop 2 (SL2) or to form long-range interactions outside the PRF region in the SuperFold predition; SL3 = Stem Loop 3; Red lines indicate pseudoknot interaction. **B)** Lower probability PRF conformation, with fully-formed SL2, colored by relative SHAPE Reactivity **C)** Dominant PRF structure prediction colored by relative Shannon entropy, labeled as in Panel A. **D)** Base-pairing probability for alternate SL2 conformation. Each dot represents a base pair in SL2. A base-pairing probability of 0.25 indicates a 25% probability of pairing for the indicated nucleotide.

The most probable, dominant structure of the PRF region, extracted from the full-length *in-vivo* secondary structure, is shown in **Fig. 3A**. In our model, the SHAPE reactivity and Shannon entropy calculation support a well-folded attenuator stem (AS) immediately upstream of the heptanucleotide slippery sequence (HSS) (**Fig 3A**; AS and HSS indicated with labeled, solid black lines). The attenuator stem has been demonstrated to be important for attenuating frameshifting in SARS-CoV(Cho et al., 2013), and previous reports suggested that the attenuator stem structure is not well conserved between SARS-CoV and SARS-CoV-2(Kelly et al., 2020). By contrast, our results suggest a SARS-CoV-2-specific fold for the attenuator stem. The highly conserved heptanucleotide slippery sequence is predicted to be single-stranded in our in-vivo structural model, which is consistent with studies on other coronaviruses(Plant et al., 2005, Plant and Dinman, 2008).

Overall, the dominant structure prediction for the H-type pseudoknot in our structural model differs from the one characterized in SARS-CoV. The H-type pseudokont in SARS-CoV is composed of three coaxially stacked stems: SL1, SL2 and the pseudoknotted helix(Plant et al., 2005). The SL1 stem, which contains the upstream pseudoknotted loop, is well folded in our consensus model as indicated by SHAPE reactivity mapping (**Fig. 3A**; labeled SL1) and Shannon entropy (**Fig. 3C**, median_Nuc(13476-13503)_ = 1.9×10^−4^; global median = 0.022). Importantly, the region reported to contain the SL2 stem(Rangan et al., 2020, Plant et al., 2005) is predicted as single-stranded in our consensus structure, and consequently does not include SL2 (**Fig 3A**; region indicated by dotted black line). Rather, the dominant structure predicted for the PRF includes a different stem, SL3, that includes the downstream pseudoknot arm (**Fig 3A**; labeled SL3). However, neither the single-stranded region nor SL3 are well-determined in our structure as indicated by Shannon entropy mapping to the region (**Fig. 3C**, median_Nuc(13503-13534)_ = 0.24; global median = 0.022, labeled with a dotted black-line and SL3, respectively).

As SuperFold calculates a partition function, lower probability base-pairing interactions are captured during structure prediction steps. We therefore checked alternative, low probability base-pair interactions captured for the PRF region. We found that the single-stranded region (**Fig. 3A**; indicated by dotted black line) forms base-pairing interaction as many as 6 different regions in the SARS-CoV-2 genome (data not shown), We sought to determine if the previously reported SL2 conformation was captured among them. Indeed, an alternate, lower probability structure containing an extended SL2 is generated in the SuperFold prediction with the attenuator stem, heptanucleotide slippery sequence, and SL1 intact. (**Fig. 3B**; alternate SL2 conformation labeled). In this variation, the SL2 stem is predicted to fold with a median probability of 20% as determined from the probabilities of each individual base-pair of the SL2 stem (**Fig. 3D;** individual base-pairs indicated with grey dots). In contrast, the SL3 stem predicted in our dominant consensus structure has as much as 80% probability of folding. The chemical probing data does not strongly support one structure over another (**Fig. 3A** and **Fig. 3B**) and likely reflects structural flexibility and pairing promiscuity for the SL2 region. Taken together, data determined *in-vivo* suggest that the frame-shifting pseudoknot in SARS-CoV-2 includes a well-folded attenuator stem, SL1, and a pseudoknot, but that the region containing the putative SL2 is conformationally flexible. Future studies are needed to explore if there is relationship between the structural flexibility and the mechanistic role of the frameshifting pseudoknot.

### The secondary structure of SARS-CoV-2 Orf1ab reveals a network of unique RNA structural elements

While the successful identification of known, functional RNA structural elements lends strong support for our methodology and for the overall secondary structural model, these known regions account for only 3% of the total nucleotide content of the SARS-CoV-2 genome; little is known about remaining 97%.

Here we report the first *in-vivo-*derived, SHAPE-constrained secondary structural model that includes a description of the base-pairing interactions for all nucleotides within a coronavirus genome (**Fig. 4A;** secondary structures described by arc plots underneath each of the three SHAPE/Shannon plots). Representative secondary structural maps of small regions extracted from the consensus prediction exemplify the types of substructures that are observed in protein-encoding regions of SARS-CoV-2 (**Fig. 4B**, structures contained within Nsp3 and spanning Nsp6&7; nt4716-5682, nt11221-12043, respectively). This resource is a valuable roadmap for ongoing studies, and to that end, a .ct file for the full-length SARS-CoV-2 genome structure is freely available (see Data Availability).

**Figure 4.**
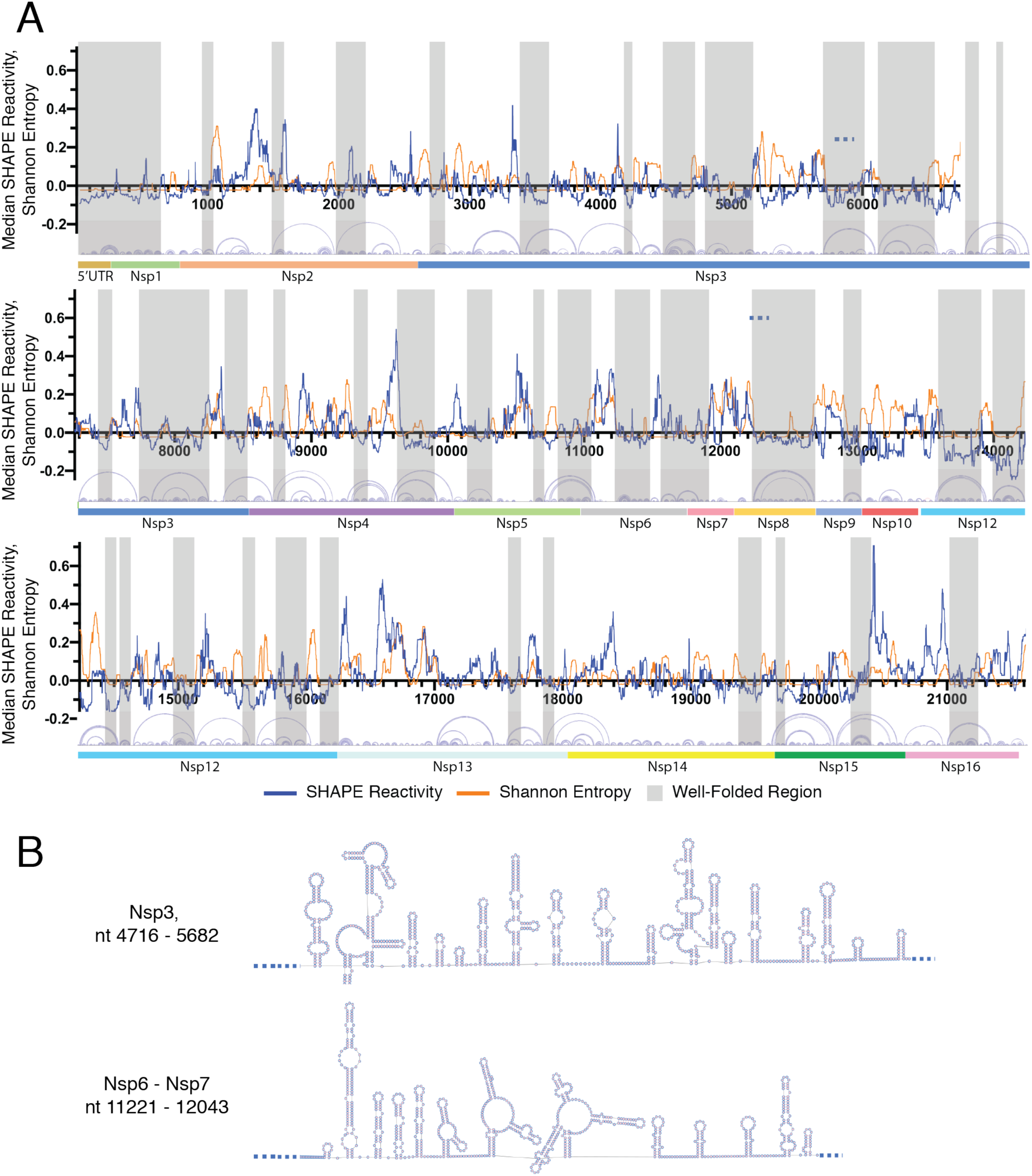
Full-length genome structure prediction of SARS-CoV-2 Orf1ab reveals a network of well-folded regions. **A)** Analysis of Shannon Entropy and SHAPE reactivities reveals 40 highly structured, well-determined domains in Orf1ab. Nucleotide coordinates are indicated on the x-axis and numbered in 1000 nucleotide intervals. Local median SHAPE reactivity and local median Shannon Entropy are indicated by blue and orange lines, respectively. Well-folded regions are shaded with grey boxes. Arc plots for all base-pairing interactions predicted by the structural model are shown beneath the local SHAPE and Shannon entropy windows, corresponding to the genomic coordinates indicated on the x-axis. The 5’UTR and non-structural protein (Nsp) domains are indicated by colored bars underneath arc plot diagrams. **B)** Representative secondary structure predictions of two regions extracted from the full-length consensus structure generated for the SARS-CoV-2 genome, with Nsp identity and genomic position indicated.

To discover additional, well-folded RNA structures within the SARS-CoV-2 genome, we used a sliding 55nt window to calculate the local median Shannon Entropy and we correlated these values with experimentally-determined SHAPE reactivities (**Fig 4A**). Only regions with both median Shannon entropy and SHAPE reactivity signals below the global median for stretches longer than 40nt, and which appear in both replicate data sets, were considered well-determined and stable. In total, we identify 40 such regions in Orf1ab (**Fig. 4B**, shaded). Hereafter, any structured region that meets these above criteria will be referred to as “well-folded.”

We also identified well-folded regions in the subgenomic RNA region (data not shown). However, our previous correlation analysis suggests that the SHAPE signal from this region includes reactivity signals from multiple RNA species, including genomic and subgenomic RNAs. Given the method deployed to construct our SHAPE-MaP sequencing libraries, it is impossible to deconvolute sgRNA data from genomic data, and new approaches will be required to separate genomic from subgenomic structures. While it will be interesting to explore this issue in subsequent studies, the following analysis focuses on stable structures within the orf1ab region, which can be uniquely determined.

To understand architectural organization of the overall “structuredness”, or base-pair content (BPC) within orf1ab, we calculated the double-strand content of individual protein domains within this region of the genome (**Fig. 5A**, grey bars). We find that all protein domains have comparable BPC, with an average of 56% (+/- 6.09%) of nucleotides involved in base-pairing interactions. However, the RNA sequences within each protein domain are not equivalently well-folded (**Fig 5A**, black bars). For example, we observe that ∼50% of nucleotides within the 5’UTR, Nsp1, Nsp6, Nsp8, and Nsp12 are concentrated in well-folded regions, suggesting these domains may be hubs for regulatory RNA structures. By contrast, Nsp13, Nsp14, and Nsp16 have <15% of their nucleotide content lies in discretely well-folded regions. At the most extreme end, Nsp10 contains no nucleotides in well-folded regions. Considering that Nsp10 is located immediately upstream of the PRF, this lack of well-folded structures may be important to direct proper frameshifting.

**Figure 5.**
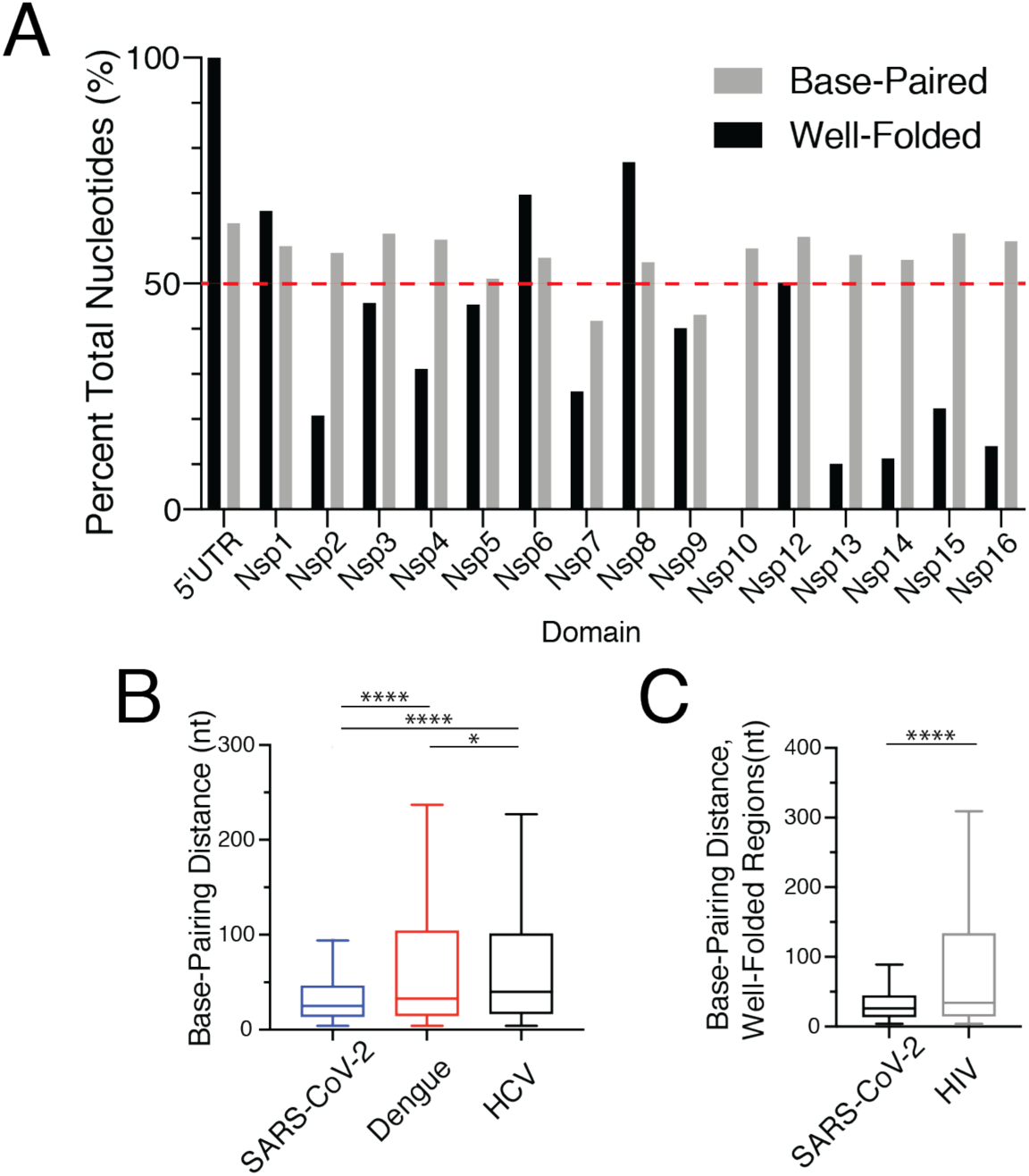
Full-length genome structure prediction of SARS-CoV-2 Orf1ab reveals a unique genome architecture. **A)** Regions encoding individual non-structural protein (Nsp) domains have comparable overall double-stranded RNA content (indicated by grey bars), but they do not adopt equally well-folded substructures (indicated by black bars). A dotted line at 50% nucleotide content has been added for clarity. **B)** SARS-CoV-2 has a shorter median base-pairing distance when compared to median base-pairing distance in previously reported, full-length genome structures for two other positive-sense RNA viruses (Mauger, *et. al.*, 2015; Dethoff, *et. al.*, 2018). Data are presented in Tukey-style box and whiskers plot as described in Fig. 1B. Asterisk definitions are below. **C)** SARS-CoV-2 has a shorter median base-pairing distance across well-folded regions of RNA when compared to those identified in HIV (Siegfried, *et al.*, 2014). Data are presented as in B). *p<0.05, ****p<0.0001 by equal variance unpaired student t test.

While analyzing the resulting secondary structural map, we noticed that the SARS-CoV-2 genome contains long-stretches of short, locally-folded stem loops (for example - **Fig. 4B**) with few long-distance base-pairing interactions such as those indicated by large arcs in typical arc plots (**Fig. 4A**). Wondering if this was quantifiable feature unique to the SARS-CoV-2 genome, we calculated the distance between base-paired nucleotides for every base-pairing interaction in our SARS-CoV-2 structural model. We compared these SARS-COV-2 base-pairing distances to those we calculated from published full-length structural models for HCV(Mauger et al., 2015) and Dengue Virus(Dethoff et al., 2018), where the data were prepared using the same structure prediction pipeline and constraints used in our study. Interestingly, the median base-pairing distance in our SARS-CoV-2 consensus model is 25nt, and is significantly smaller than the median base-pairing distance in the HCV (median=40nt) and Dengue Virus (median=33nt) consensus models (**Fig. 5B**). Even more, the upper bound of the interquartile range (IQR) that describes the distribution of base-pairing distances in Dengue and HCV genomes is much higher than the same bound in the SARS-CoV-2 genome (SARS-CoV-2 75^th^ percentile = 46nt; Dengue Virus 75^th^ Percentile = 104nt; HCV 75^th^ percentile = 101nt). This suggests SARS-CoV-2 has fewer long-distance base-paring interactions compared to Dengue and HCV genome.

We also calculated the median base-pairing distance for the well-folded regions of the SARS-CoV-2 genome and compared the result to well-folded regions previously identified using the same Low Shannon/Low SHAPE signatures in the HIV genome(Siegfried et al., 2014). We found that although there is no significant difference in the size of well-folded regions in the SARS-CoV2 and HIV genomes (data not shown), the median base-pairing distance in the well-folded regions of SARS-CoV-2 (median = 26nt) is significantly lower than the base-pairing distance in well-folded regions of HIV (median = 34nt) (**Fig. 5C).** Similarly, the upper bound of the IQR that describes the distribution of base-pairing distances in well-folded regions of the HIV genome is much higher than the same bound in SARS-CoV-2 (SARS-CoV-2 75^th^ percentile = 44nt; HIV 75^th^ percentile = 133nt).

Taken together, these results suggest that the SARS-CoV-2 genome folds into more local secondary structures, such as the short stem-loops in **Fig. 4B**, and contains fewer long-range base-pairing interactions than observed for other positive-sense RNA viruses. Given the exceptional size of the coronavirus genome (∼30kb) relative to those of the positive-sense RNA viruses compared here (∼10kb), it is possible that the short base-pairing distance of SARS-CoV2 may carry functional implications for maintaining genomic stability, preserving fidelity of translation, and evading innate immune response.

### The overall structured-ness of the SARS-CoV-2 is conserved across all β-coronaviruses

Synonymous mutations rates have been used previously to lend evolutionary support for well-folded RNA secondary structures in other positive-sense RNA viruses(Dethoff et al., 2018, Tuplin et al., 2002, Assis, 2014, Simmonds and Smith, 1999). This body of work has suggested lower synonymous rates for double-stranded nucleotides when compared to single-stranded nucleotides in viral RNAs, likely reflecting an evolutionary pressure to maintain base-pairing interactions of double-stranded nucleotides. We therefore computed relative synonymous mutation rates to determine how evolutionary pressure is applied to single- and double-stranded regions of the SARS-CoV2 genome.

To generate the codon-based alignment, 33 full-length genome sequences from the NCBI Taxonomy database(Benson et al., 2018) were collected, including 12 SARS-CoV genomes, 8 MERS-CoV genomes, and 13 more distantly related β-coronaviruses genomes. This alignment was then used to calculate synonymous and non-synonymous mutation rates (dS and dN, respectively) for each codon in the SARS-CoV2 orf1ab region using A Fast, Unconstrained Bayesian AppRoximation for inferring selection (FUBAR) (Murrell et al., 2013). We then separated dS and dN into single- or double-stranded bins as predicted in our consensus model. The strandedness of each codon was determined by the strandedness at the third position of the codon(Dethoff et al., 2018).

In the “All β-Coronavirus” alignment, we observed a significantly lower synonymous mutation rate (p<0.0001) for double-stranded codons (median = 3.765; IQR = 3.034-4.978) when compared to single-stranded codons (median = 4.189; IQR = 3.232 - 5.562) in our consensus model (**Fig. 6A**). In contrast, there was no significant difference (p = 0.86) observed for non-synonymous mutation rates (dN) at single- (median = 0.4535; IQR = 0.103 – 0.685) or double-stranded codons (median = 0.453; IQR = 0.139 – 0.675) (**Fig. 6B**) as dN reflects changes at the amino acid level. This suggests that double-stranded regions of the SARS-CoV-2 genome experience stronger selective pressure against synonymous mutations than single-stranded regions, which lends support to our consensus model and suggests evolutionary maintenance of the observed secondary structure. Because an all β-coronavirus alignment was used, our results indicate that the structural organization and overall base-pairing content of Orf1ab is a conserved feature of the β-coronavirus family.

**Figure 6.**
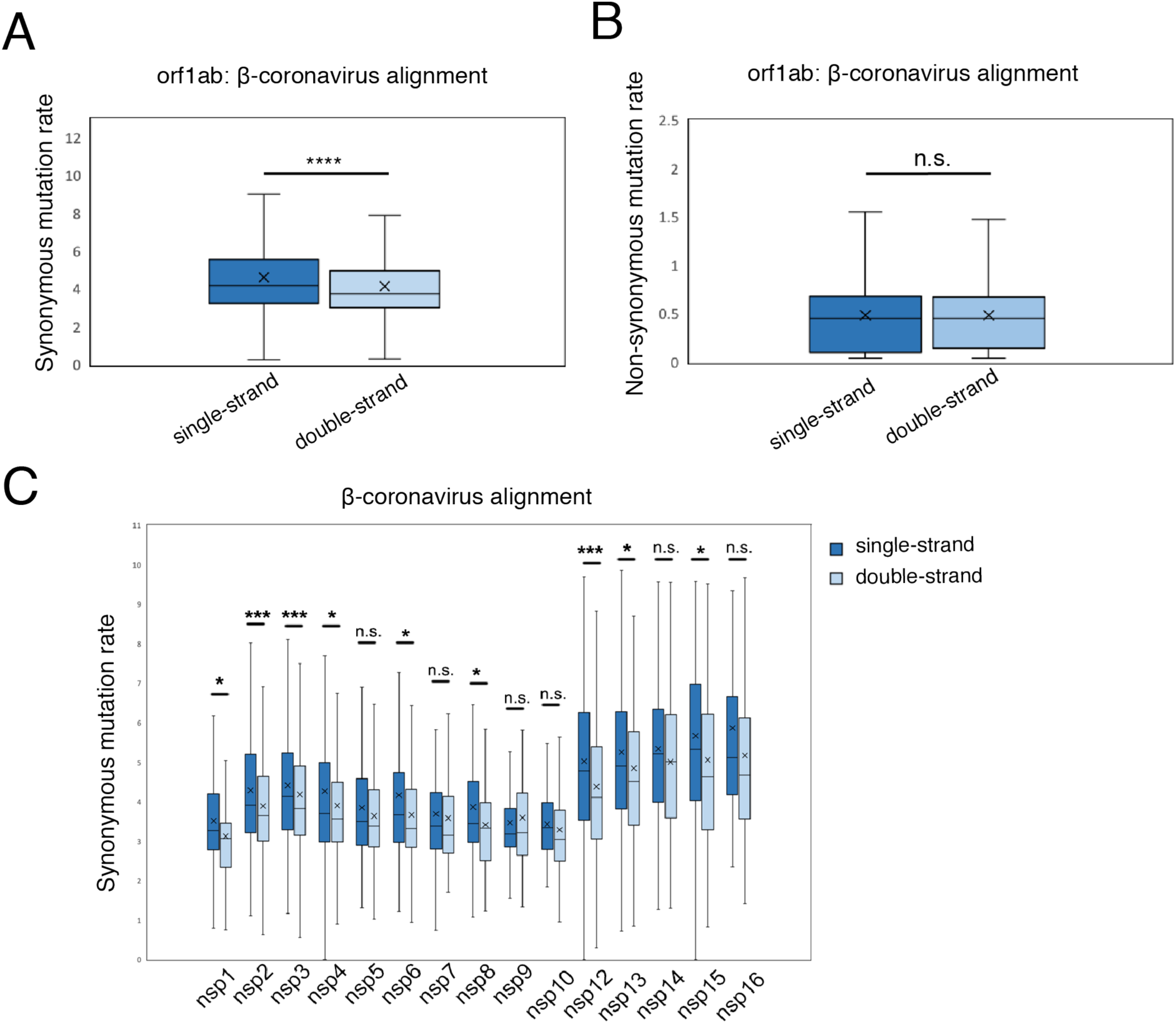
Structure-dependent variations in synonymous mutation rates suggest that all β-coronaviruses have highly structured genomes (high BPC). **A)** Synonymous mutation rates calculated across all β-coronaviruses for single- and double-stranded nucleotides of Orf1ab. Data are presented in Tukey-style box and whiskers plot as described in Fig. 1B. **B)** Non-synonymous mutation rates calculated across all β-coronaviruses for single- and double-stranded nucleotides of Orf1ab. Data are presented as in (A). **C)** Comparison of synonymous mutation rates for single- and double-stranded nucleotides within individual protein domains, calculated across all β-coronaviruses. Data are presented as in (A). n.s. not significant,*p<0.05, ***p<0.001 ****p<0.0001 by equal variance unpaired student t test.

When analyzing relative synonymous mutation rates within individual protein domains, we observed significantly decreased synonymous mutation rates for double-stranded codons in Nsp1, Nsp2, Nsp3, Nsp4, Nsp6, Nsp8, Nsp12, Nsp13, and Nsp15 (**Fig. 6C**). Consistent with this, Nsp1, Nsp6, Nsp8, and Nsp12 have >50% of their nucleotides localized within well-folded regions (**Fig. 5A**, black bars). Taken together, this suggests that certain protein-coding domains contain regions of RNA secondary structure that are conserved across β-Coronaviruses. For example, Nsp8, which is the most well folded domain in SARS-CoV-2, is likely well-folded in other β-Coronaviruses.

By contrast, the base pairing content of Nsp5, Nsp7, Nsp9, Nsp10, Nsp14, and Nsp16 does not appear to be conserved, as there is no significant difference in synonymous mutation rates of single- and double-stranded codons (**Fig. 6C**). Consistent with this, Nsp14 and Nsp16 were shown to have <15% of their nucleotides in well-folded regions, while Nsp10 does not contain any well-folded nucleotides (**Fig. 5A**). Not only does this analysis support the observation that these regions of RNA are not well-folded in SARS-CoV-2, our data suggest these regions may not be well folded in other β-Coronaviruses.

### Evolutionary analysis for individual well-folded regions of the SARS-CoV-2 genome identifies several conserved regions

To further prioritize structural elements that may have conserved functional roles in the SARS-CoV2 life cycle, we next applied our synonymous mutation rate analysis to each of the 40 discrete well-folded domains identified by Low Shannon/Low SHAPE signatures (**Fig. 4B, Table S1**). Four regions (regions 23, 25, 34, and 36) that are well-determined based on their low median Shannon Entropy values (**Table S1**) and SHAPE reactivity data showed significantly decreased synonymous mutation rates at double-stranded codons when compared to single-stranded codons across the β-coronavirus alignment (**Fig. 7A, 7B)**. Among those structures, region 25 and 34 are found at protein domain boundaries. Region 25 ends exactly at the Nsp8/9 domain boundary, while Region 34 spans the Nsp12/13 boundary. Region 23, 34, and 36 (**Fig. 7C, Fig. 7E, Fig. 7F**) contain a series of stem-loops with small bulges. Region 25 contains a long-range duplex that closes a clover-leaf like structure with 8 stem-loops radiating from a central loop (**Fig. 7D**). This hub, or multi-helix junction might represent a promising drug target, as multi-helix junctions often contain binding pockets with high binding affinity and selectivity for small molecules(Warner et al., 2018).

**Figure 7.**
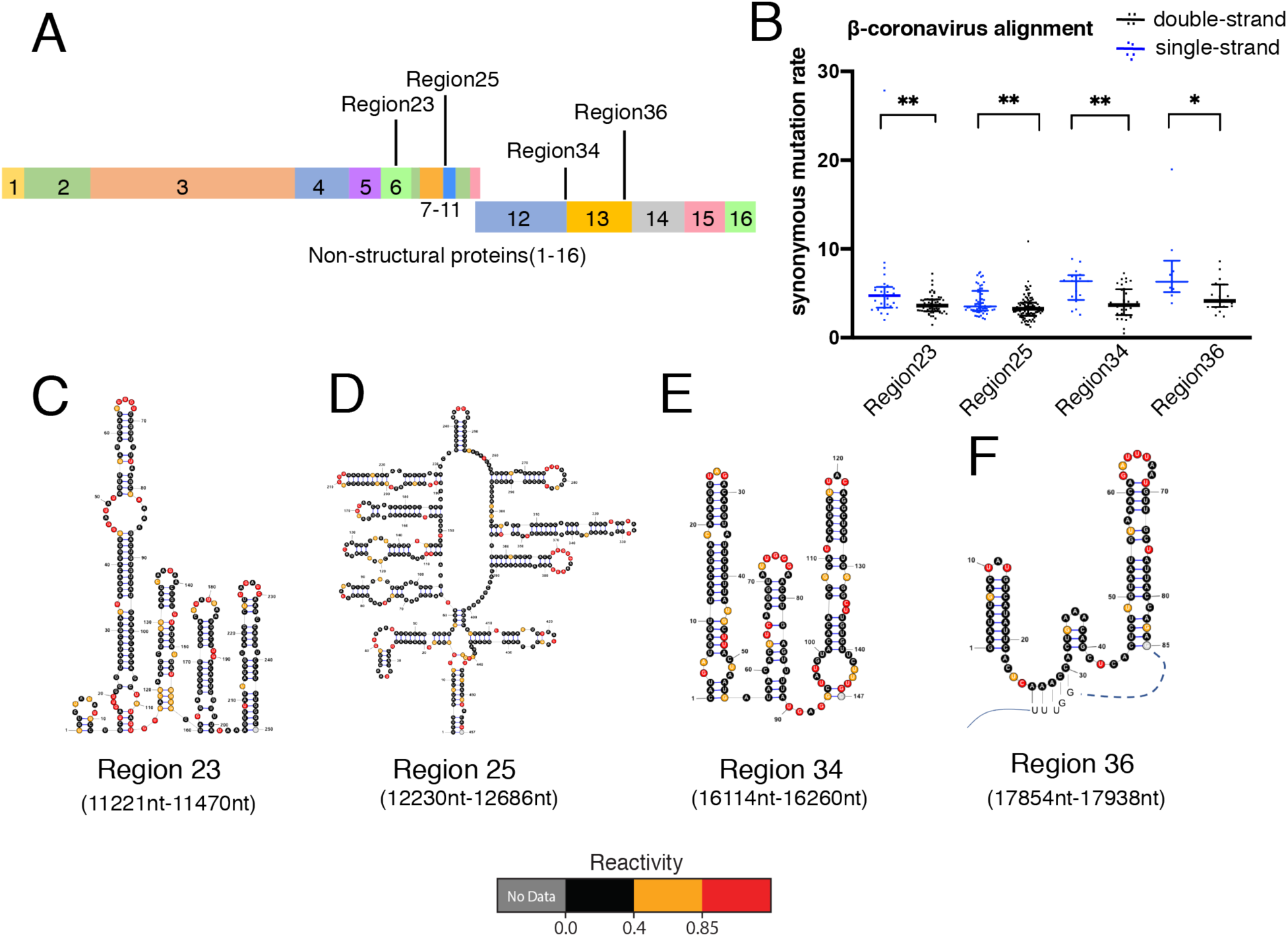
Analysis of synonymous mutation rates within individual well-folded regions of the SARS-CoV-2 genome identifies four regions that appear to be conserved across β-coronaviruses. **A)** Schematic of well-folded regions in SARS-COV2 genome supported by Synonymous mutation rate analysis in β-coronaviruses. **B)** Synonymous mutation rate separated by stranded-ness in four individual well-folded regions. Data are plotted with a line indicating the median, and whiskers indicating the interquartile range central. *p<0.05, **p<0.01 by equal variance unpaired student t test. **C), D), E), F)** RNA secondary structure diagrams of four well-folded regions supported by analysis of synonymous mutation rates, colored by SHAPE reactivities, with genomic coordinates indicated below and in (A).

Within the Sarbecovirus subgenus, we were able to identify four regions (regions 15, 22, 24, 27, and 30) that are well-determined in our secondary structural model (based on low median Shannon Entropy (**Table S1**) and SHAPE reactivity data) with significantly decreased synonymous mutation rates in double-stranded relative to single-stranded codons (**Fig. 8A, 8B**). Among these structures, Region 24 contains two discrete multi-helix junctions, each with at least three stems radiating from large central loops (**Fig. 8C**). Region 27, which contains a series of six stem-loops, is particularly significant because it is only 100nt downstream of the PRF in Nsp12 (**Fig 8D)**. Region 15, like Region 24, contains several well-determined long-range duplexes that segment the region into two discrete multi-helix junctions (**Fig. 8E**). Region 22 contains a series of well-folded loops and it spans the Nsp5/6 boundary (**Fig. 8F**). Region 30 is a single stem-loop with bulges that divide the stem into distinct duplexes (**Fig. 8G**)

**Figure 8.**
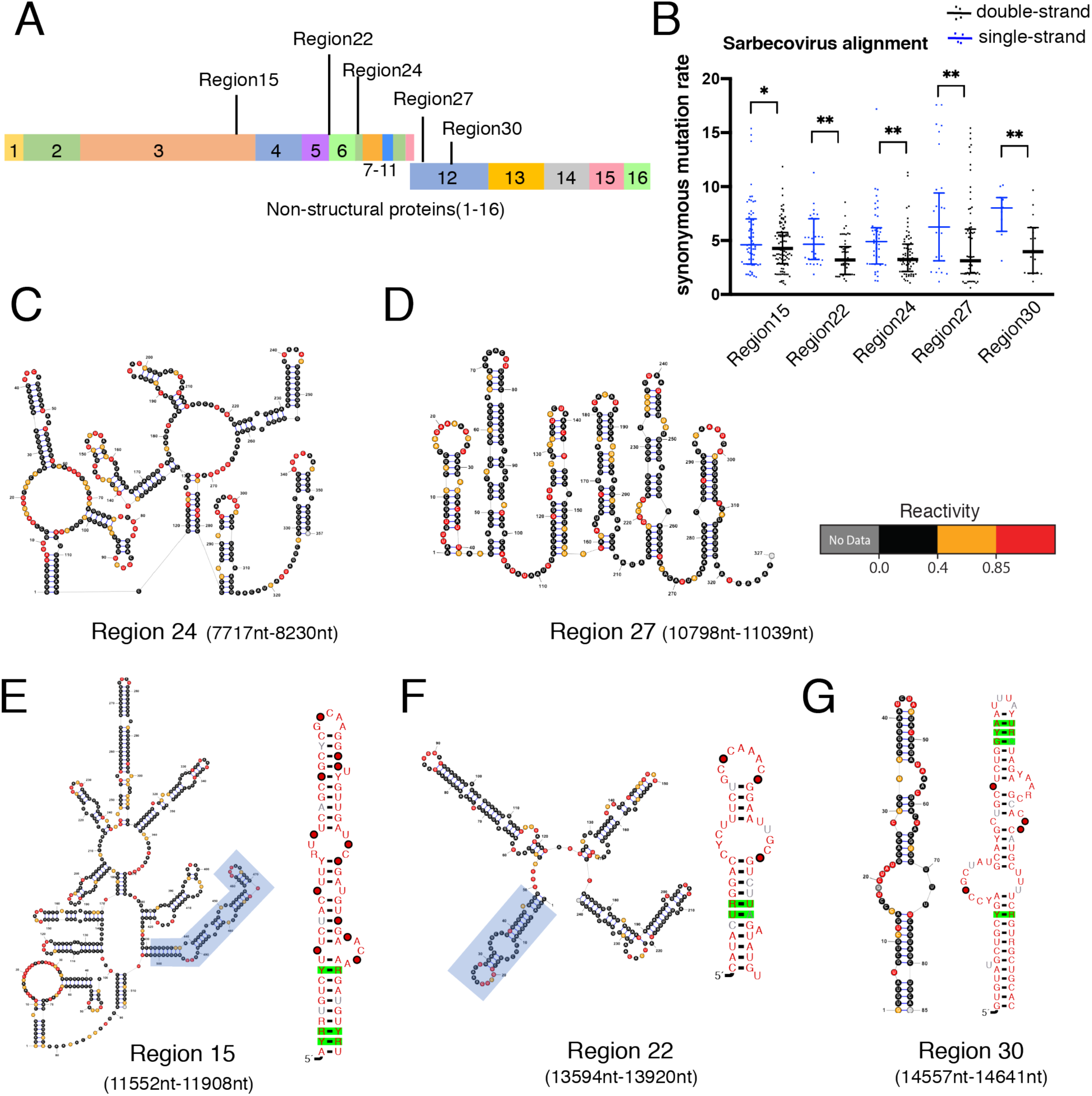
Analysis of synonymous mutation rates and covariation within individual regions of the SARS-CoV-2 genome pinpoints five regions that are conserved only within the sarbecovirus subgenus. **A)** Schematic of well-folded regions in the SARS-COV2 genome supported by Synonymous mutation rate analysis in the sarbecovirus subgenus. **B)** Synonymous mutation rate separated by stranded-ness in five individual well-folded regions. Data are plotted with a line indicating the median, and whiskers indicating the interquartile range central. *p<0.05, **p<0.01 by equal variance unpaired student t test. **C)**,**D)** RNA secondary structures of two well-folded regions colored by SHAPE reactivity **E), F), G)** RNA secondary structure diagrams of three well-folded regions supported by both synonymous mutation rate analysis and covariation in sarbecoviruses, colored by SHAPE reactivities. Green boxes indicate significant covariation base pairs tested by Rscape-RAFSp(e-value<0.05). Consensus nucleotides are colored by relative degree of sequence conservation within the alignment (75% identity in gray, 90% identify in black, 97% identity in red). Individual nucleotides are represented by circles according to their positional conservation and percentage occupancy thresholds (50% occupancy in white, 75% occupancy in grey, 90% occupancy in black, 97% occupancy in red). Multiple sequence alignment files are provided in supplementary materials.

To look for evolutionary evidence that directly supports conservation of specific base-pairing interactions and secondary structures, we performed covariation analysis on the 5 structures that are supported by Sarbecovirus-specific synonymous mutation rates. We visualized the base-pair covariation levels using R-chie (Lai et al., 2012) and we used R-scape version 0.2.1(Rivas et al., 2017) with the RAFSp statistics(Tavares et al., 2019) to test the statistical significance of putatively covarying base-pairs. Specifically, we identified 3 regions (15, 22, and 30) that have covariation support **(Fig. 8E-G)**. Region 15 has three significantly covarying base-pairs at the terminus of the downstream stem loop (e-value < 0.05, **Fig. 8E**); Region 22 has 2 nucleotides with have one-sided variation in the most upstream stem loop (e-value < 0.05, **Fig. 8F**); Region 30 has 3 covarying base-pairs at the very top of the stem loop, and a single covarying pair at the bottom portion of the stem (e-value < 0.05, **Fig. 8G**). Taken together, these results suggest the existence of stable, evolutionarily conserved structural elements that merit subsequent functional analysis.

## Discussion

Here we establish that the SARS-CoV-2 genomic RNA has an extraordinarily complex molecular architecture, filled with elaborate secondary and tertiary structural features that persist in-vivo and which are conserved through time, suggesting that this network of RNA secondary structural elements plays a functional role in the virus lifecycle. Indeed, the SARS-CoV-2 genome contains more well-determined RNA structures than any virus studied to date, suggesting that its inherent “structuredness” contributes in a unique way to viral fitness. This RNA secondary structural complexity is not just confined to untranslated regions of the genome, as protein-coding sections of the SARS-CoV-2 open reading frame are among the most well-structured regions. Thus, as observed for HCV, coronavirus reading frames experience evolutionary pressure that simultaneously shapes both protein sequence and the surrounding RNA structures in which the proteins are encoded (a “code within the code”)(Pirakitikulr et al., 2016). The secondary structure that we report is well-determined based on available metrics in the field(Siegfried et al., 2014). It is both a roadmap for navigating the vast RNA landscape in coronaviruses, and a resource for orthogonal studies by others. As such, the data reported here are all publicly available for analysis and comparison by others https://github.com/pylelab/SARS-CoV-2_SHAPE_MaP_structure.

Well-determined secondary structures of long RNA molecules are typically difficult to obtain *in-vivo*(Mitchell et al., 2019, Leamy et al., 2016). They are usually derived from transcripts that have been refolded and probed in-vitro, or from isolated cellular transcripts that have been stripped of cellular components(Smola et al., 2015a, Siegfried et al., 2014). What is particularly surprising about this SARS-CoV-2 study, and the high quality of the resulting secondary structure, is the fact that it was entirely determined *in-vivo*, using infected cells that were treated directly with chemical probes. This may be attributable to the fact that SARS-CoV-2 genomic RNA is so abundant in the infected cell, ultimately becoming ∼65% of the total cellular RNA(Kim et al., 2020). With so much RNA material, it becomes possible to maximize the signal to noise ratio in chemical probing experiments. In addition, the abundance of SARS-CoV-2 RNA may overwhelm the cell’s ability to coat transcripts with nonspecific RNA binding proteins, which can otherwise limit accessibility of chemical probes. That said, it will be interesting to compare the structure reported here with that obtained “*ex vivo*” (stripped of protein), as that ΔSHAPE approach provides a useful way to flag possible protein binding sites(Smola et al., 2015a).

The resulting experimental secondary structure provides new insights into known coronaviral RNA motifs, and leads to the prediction of new ones that are likely to regulate viral function. The near perfect structural homology of motifs at the 5’ terminus for SARS-CoV-2 and other β-coronavirus genomes suggests that the function of these upstream elements is conserved in coronaviruses (reviewed in (Yang and Leibowitz, 2015)). Furthermore, because our SARS-CoV-2 secondary structure was determined *in-vivo*, our findings validate previous coronavirus structural models of 5’-elements, as our data were obtained in a biologically relevant context.

Our SARS-CoV-2 secondary structure at the 3’ viral terminus largely agrees with previous studies on other β-coronavirus genomes (reviewed in(Yang and Leibowitz, 2015)). However, our model of the 3’ viral terminus deviates in one important way. Neither the raw SHAPE reactivity data nor the subsequent secondary structure prediction supports formation of a pseudoknot proposed between the base of the BSL and SLI. Indeed, the putative pseudoknot conformation is mutually exclusive with the well-structured stem that we report at the base of the BSL. However, both conformations are proposed to be essential in MHV(Goebel et al., 2004), so it is possible that the pseudoknot exists as a minority conformation, or is transiently folded in SARS-CoV-2. Alternatively, because our structure represents the first detailed description of a coronavirus 3’UTR structure *in-vivo*, it is possible this pseudoknot is not present in other viruses.

Arguably the best-studied structural element in coronaviruses is the programmed ribosomal frameshifting pseudoknot (PRF). Required for proper replicase translation in all coronavirus family members, the PRF adopts different conformations in the various coronaviruses, including three-stemmed, two-stemmed, and kissing-loop pseudoknots (Baranov et al., 2005, Plant and Dinman, 2008). The core of the SARS-CoV PRF, which shares an almost identical sequence with SARS-CoV-2, is predicted to form a three-stem pseudoknot comprised of SLI, SL2, and a pseudoknot helix, with an additional upstream attenuator stem that is poorly conserved in SARS-CoV-2(Kelly et al., 2020). Our SHAPE reactivity and structure prediction are consistent with the existence of an attenuator stem, SL1, and the pseudoknot. However, our data indicate that the region corresponding to SL2 is conformationally flexible, adopting an SL2 stem with only a 20% probability. Consistent with our reported distribution of structural isoforms, Kelly et al. use a reporter assay to suggest that the frequency of successful frameshifting in SARS-CoV-2 is about 20%(Kelly et al., 2020), indicating that the observed conformational variability of SL2 may be functional. Indeed, SL2 might function like a switch: When SL2 is formed (∼20% of the time), frameshifting occurs. When unfolded or forming base-pairs with structures outside the PRF region, frameshifting would not occur. Further studies are therefore required to explore the relationship between SL2 formation and SARS-CoV-2 frame-shifting efficiency.

The study reported here provides a structure prediction for every single nucleotide in the SARS-CoV-2 genome, enabling us to simultaneously interrogate both global and local features of genome architecture. One can make two major observations about the global architecture the SARS-CoV-2 genome. First, this *in-vivo* derived, SHAPE-constrained model strongly agrees with the high double-strand RNA content predicted from the entirely *in silico* model recently reported by our lab (Tavares et al., 2020). Because the data herein were obtained *in-vivo*, this work confirms that the unusually high double-strand content is maintained in a cellular context. Secondly, analysis of the experimental secondary structure reveals that the SARS-CoV-2 genome has a shorter median base-pairing distance when compared with other positive-sense RNA viral genomes, suggesting a role for extreme compaction in the function of coronaviral genomes. Downstream analysis of synonymous mutation rates suggests that global architectural features are conserved across β-coronaviruses. Considering the exceptional size of these genomes, the high degree of dsRNA content may represent an evolutionary strategy to enhance genome stability, as duplex RNA undergoes self-hydrolysis at a much slower rate than single-stranded RNA and it is more resistant to cellular nucleases(Regulski and Breaker, 2008, Wan et al., 2011). Interestingly, single-stranded regions in mRNA have been shown to mediate phase separation at high cellular RNA concentrations(Van Treeck et al., 2018). Because SARS-CoV-2 RNA is very abundant *in-vivo* (up to 65% of total cellular RNA content (Kim et al., 2020)) it is possible the high dsRNA content may provide a strategy to avoid phase separation during infection. The preference for abundant locally folded, short stem-loop structures in β-coronavirus genomes may also provide a conserved strategy for innate immune evasion. Pattern recognition receptors such as MDA5(Dias Junior et al., 2019) and ADAR modification(Nishikura, 2010) proteins recognize long RNA duplexes as part of host defense processes, which could obviously be avoided by keeping duplex lengths short.

Analysis of local features within the genome pinpoints 40 well-folded regions within the SARS-CoV-2 orf1ab region. Of these 40 regions, at least five are conserved across all β-coronaviruses and four are sarbecovirus specific. Four of the nine regions (Region 25, Region 34, Region 22, Region 24) span boundaries between non-structural proteins, which may have relevance for polyprotein translation. Previous studies have shown that RNA secondary structures can slow the rate of ribosome translocation(Chen et al., 2013) and ribosome stalling is known to be important for proper folding of nascent polypeptides(Collart and Weiss, 2020). Conserved, well-folded RNA structures at protein domain boundaries may therefore slow or stall translocating ribosomes, thus allowing individual non-structural proteins in the large Orf1a and Orf1ab poly-proteins to fold into their native conformations.

Intriguingly, three of the nine well-folded regions (Region 15, Region 24, Region 25) contain complex, multi-helix junctions, or structural hubs. This is significant because multi-helix junctions often comprise the core of RNA tertiary structures, like group II self-splicing introns, riboswitches and other regulatory elements. Because these elements are likely to contain well-defined pockets, they often bind specifically to small molecules, and therefore serve as possible drug targets (Warner et al., 2018, Hewitt et al., 2019, Fedorova et al., 2018).

One important cautionary observation from our work is the poor correlation of SHAPE reactivities between two *in-vivo* biological replicates for regions encoding the subgenomic RNAs. Previous *in silico* work from our lab has shown that individual subgenomic RNAs (sgRNAs), such as the N sgRNA, fold differently than the corresponding regions in the genomic RNA due to differences in upstream sequence context(Tavares et al., 2020). Though our tiled-amplicon design affords sequencing coverage for the entire SARS-CoV-2 genome, it precludes deconvolution of reactivity signals for regions shared between genomic- and subgenomic RNAs. This underscores the need for methodological innovations that accurately assess the structural content specific to subgenomic RNA molecules. Absent such methodological advances, we caution others when interpreting reactivities from the subgenomic region.

The *in-vivo*-determined SARS-CoV2 secondary structure present here provides a roadmap for functional studies of the SARS-CoV2 genome and insights into mechanisms of the SARS-CoV-2 life cycle. Evolutionary support for consensus model across β-coronaviruses hints at conserved strategies for genome stability, translation fidelity, and innate immune evasion. Finally, the identification of individual well-folded regions conserved across β-coronaviruses, and within the sarbecovirus subgenus, provide potential targets for the study of regulatory elements, and the search for much-needed therapeutically active small molecules.

## Acknowledgments

We would like to thank Dr. Li-Tao Guo for preparing and sharing MarathonRT enzyme, and Dr. Ananth Kumar and Gandhar Mahadeshwar (Pyle lab, Yale University) for thoughtful comments on the manuscript. We would also like to thank Dr. Mark Boerneke (Weeks Lab, UNC Chapel Hill) for providing data upon request. This work was supported by the Howard Hughes Medical Institute; the National Institutes of Health (R01 HG009622 to A.M.P.); the NIH Grant T32AI055403 (to N.C.H.); China Scholarship Council (CSC)-Yale World Scholars Program in Biomedical Sciences (to H.W.); Funding for open access charge was provided by the Howard Hughes Medical Institute.

## Author Contributions

N.C.H., H.W., and C.W. conducted experiments. N.C.H., H.W., C.W., and A.M.P. designed experiments. N.C.H., H.W., R.C.A.T, A.M.P. wrote the paper.

