## Supplemental Tables S1-S3 for "Comprehensive in-vivo secondary structure of the SARS-CoV-2 genome reveals novel regulatory motifs and mechanisms"

**Table S1.** Well-determined region in Orf1ab region

| Region | Window | Start | End | Size | Protein Domain | Median Shannon Entropy |
|---|---|---|---|---|---|---|
| 1 | 1 | 1 | 622 | 622 | 5'UTR, Nsp1 | 7.69E-05 |
| 2 | 1 | 944 | 1026 | 83 | Nsp2 | 1.75E-06 |
| 3 | 1 | 1478 | 1572 | 95 | Nsp2 | 2.62E-02 |
| 4 | 1 | 1968 | 2188 | 221 | Nsp2 | 5.07E-04 |
| 5 | 1 | 2682 | 2800 | 119 | Nsp3 | 9.19E-03 |
| 6 | 1 | 3416 | 3597 | 182 | Nsp3 | 5.24E-05 |
| 7 | 1 | 4169 | 4232 | 64 | Nsp3 | 3.01E-06 |
| 8 | 1 | 4471 | 4713 | 243 | Nsp3 | 4.82E-04 |
| 9 | 1 | 4791 | 5162 | 372 | Nsp3 | 4.55E-04 |
| 10 | 1 | 5693 | 6013 | 321 | Nsp3 | 2.07E-03 |
| 11 | 1 | 6116 | 6549 | 434 | Nsp3 | 4.24E-04 |
| 12 | 1 | 6786 | 6887 | 102 | Nsp3 | 4.88E-03 |
| 13 | 1 | 7025 | 7072 | 48 | Nsp3 | 1.06E-03 |
| 14 | 2 | 7413 | 7518 | 106 | Nsp3 | 1.55E-04 |
| 15 | 2 | 7717 | 8230 | 514 | Nsp3 | 1.29E-03 |
| 16 | 2 | 8350 | 8512 | 163 | Nsp3 | 4.90E-04 |
| 17 | 2 | 8702 | 8789 | 88 | Nsp4 | 6.45E-04 |
| 18 | 2 | 9295 | 9398 | 104 | Nsp4 | 2.76E-02 |
| 19 | 2 | 9612 | 9886 | 275 | Nsp4 | 1.70E-02 |
| 20 | 2 | 10129 | 10312 | 184 | Nsp5 | 3.00E-04 |
| 21 | 2 | 10630 | 10687 | 58 | Nsp5 | 3.53E-04 |
| 22 | 2 | 10798 | 11039 | 242 | Nsp5, Nsp6 | 7.67E-04 |
| 23 | 2 | 11221 | 11470 | 250 | Nsp6 | 2.34E-03 |
| 24 | 2 | 11552 | 11908 | 357 | Nsp6, Nsp7 | 2.09E-03 |
| 25 | 2 | 12230 | 12686 | 457 | Nsp8 | 1.86E-03 |
| 26 | 2 | 12895 | 13030 | 136 | Nsp9 | 1.55E-02 |
| 27 | 2 | 13594 | 13920 | 327 | Nsp12 | 7.39E-04 |
| 28 | 2 | 13993 | 14230 | 238 | Nsp12 | 1.19E-02 |
| 29 | 3 | 14444 | 14532 | 89 | Nsp12 | 4.27E-04 |
| 30 | 3 | 14557 | 14641 | 85 | Nsp12 | 1.05E-05 |
| 31 | 3 | 14973 | 15136 | 164 | Nsp12 | 9.07E-04 |
| 32 | 3 | 15510 | 15608 | 99 | Nsp12 | 1.59E-04 |
| 33 | 3 | 15767 | 16005 | 239 | Nsp12 | 4.29E-04 |
| 34 | 3 | 16114 | 16260 | 147 | Nsp12 | 2.69E-03 |
| 35 | 3 | 17580 | 17677 | 98 | Nsp13 | 1.64E-04 |
| 36 | 3 | 17854 | 17938 | 85 | Nsp13 | 6.09E-04 |
| 37 | 3 | 19373 | 19550 | 178 | Nsp14 | 1.52E-02 |
| 38 | 3 | 19665 | 19735 | 71 | Nsp15 | 2.32E-05 |
| 39 | 3 | 20248 | 20408 | 161 | Nsp15 | 7.72E-03 |
| 40 | 3 | 20668 | 20792 | 125 | Nsp16 | 4.91E-06 |

**Table S2.** Gene-specific RT primers

| Primer Name | Sequence |
|---|---|
| RT_SC2_Amplicon_1 | TTTTTTTTTGTCATTCTCC |
| RT_SC2_Amplicon_2 | ATGTTGAGTACATGACTGT |
| RT_SC2_Amplicon_3 | TAACATGTTCAACACCAGT |
| RT_SC2_Amplicon_4 | AATCATTTCATCTGTGAGC |
| RT_SC2_Amplicon_5 | TAATACCTATTGGCAAATC |
| RT_SC2_Amplicon_6 | AACCACCTAACTGACTATG |
| RT_SC2_Amplicon_7 | TAACTCTGGAAAAATCTGT |
| RT_SC2_Amplicon_8 | TAACATTATCGCTACCAAC |
| RT_SC2_Amplicon_9 | TTAGTAAGTGCAGCTACTG |
| RT_SC2_Amplicon_10 | AAGCAGTTTGTGTAGTACC |
| RT_SC2_Amplicon_11 | TATCTAAAACGGCAATTCC |
| RT_SC2_Amplicon_12 | TACCAACTGCACTAAAAAC |
| RT_SC2_Amplicon_13 | AATTAGACATTAAAACACC |
| RT_SC2_Amplicon_14 | AAACATAAAATGTTTTACC |
| RT_SC2_Amplicon_15 | TTTGTTGACTATCATCATC |
| RT_SC2_Amplicon_16 | TTAGTCAAATTCTCAGTGC |

**Table S3.** Gene-specific PCR Primers

| | |
|---|---|
| R_PCR_SC2_Amplicon_1 | TTTTTTGTCATTCTCCTAAGAAG |
| R_PCR_SC2_Amplicon_2 | TTGAGTACATGACTGTAAACTACAT |
| R_PCR_SC2_Amplicon_3 | CATGTTCAACACCAGTGTCTGTA |
| R_PCR_SC2_Amplicon_4 | CATTTCATCTGTGAGCAAAG |
| R_PCR_SC2_Amplicon_5 | TACCTATTGGCAAATCTACCAAT |
| R_PCR_SC2_Amplicon_6 | CACCTAACTGACTATGACTAAAA |
| R_PCR_SC2_Amplicon_7 | CTCTGGAAAAATCTGTATTATTAGG |
| R_PCR_SC2_Amplicon_8 | CATTATCGCTACCAACACATGTA |
| R_PCR_SC2_Amplicon_9 | GTAAGTGCAGCTACTGAAAAGCA |
| R_PCR_SC2_Amplicon_10 | CAGTTTGTGTAGTACCGGCA |
| R_PCR_SC2_Amplicon_11 | CTAAAACGGCAATTCCAGTT |
| R_PCR_SC2_Amplicon_12 | CCAACTGCACTAAAAACTCTAGG |
| R_PCR_SC2_Amplicon_13 | TTAGACATTAAAACACCTAAAGC |
| R_PCR_SC2_Amplicon_14 | ACATAAAATGTTTTACCTTCATG |
| R_PCR_SC2_Amplicon_15 | TGTTGACTATCATCATCTAACCA |
| R_PCR_SC2_Amplicon_16 | AGTCAAATTCTCAGTGCCACAA |
| F_PCR_SC2_Amplicon_1 | GTCACGCCTAAACGAACATG |
| F_PCR_SC2_Amplicon_2 | TCTGGAGTAAAAGACTGTGTTGT |
| F_PCR_SC2_Amplicon_3 | GTGATTGCCTTGGTGATATT |
| F_PCR_SC2_Amplicon_4 | TATATTCTAAGCACACGCCTATT |
| F_PCR_SC2_Amplicon_5 | TTAGAATTAGCTATGGATGAATT |
| F_PCR_SC2_Amplicon_6 | ACAGCTAGGTTTTTCTACAGGTG |
| F_PCR_SC2_Amplicon_7 | TTATTGTAAATCACATAAACCAC |
| F_PCR_SC2_Amplicon_8 | TAAGGAATTACTTGTGTATGCTG |
| F_PCR_SC2_Amplicon_9 | TGTAACAGCTTTAAGGGCCAATT |
| F_PCR_SC2_Amplicon_10 | TGTGGCTATGAAGTACAATTATG |
| F_PCR_SC2_Amplicon_11 | AAGAGAAGTGGGTTTTGTCG |
| F_PCR_SC2_Amplicon_12 | ATAAATATTATAATTTGGTTTTTACTATTA |
| F_PCR_SC2_Amplicon_13 | TATGGACAACAGTTTGGTCCAAC |
| F_PCR_SC2_Amplicon_14 | GATTACCAAGGTAAACCTTTGGA |
| F_PCR_SC2_Amplicon_15 | CTCATGAAGTGTGATCATTGTGG |
| F_PCR_SC2_Amplicon_16 | ATTAAAGGTTTATACCTTCCCAG |
